# Efficient memory sampling by hippocampal attractor dynamics with intrinsic oscillation

**DOI:** 10.64898/2026.03.05.709774

**Authors:** Tatsuya Haga

## Abstract

Hippocampus is known to replay activity patterns to recall and process memories, which is often related to Hopfield-type attractor dynamics. Another line of theoretical studies suggests that hippocampal replay prioritizes replay of experiences to accelerate value learning for efficient decision making. It is unknown how hippocampal attractor dynamics perform prioritized memory sampling, and more broadly, how we can consistently relate dynamical (bottom-up) and functional (top-down) theories of hippocampal replay. In this paper, we propose an extended Hopfield-type attractor network model with momentum, kinetic energy, and conservation of the total energy, which is called momentum Hopfield model. We show that our model can be interpreted as CA3-CA1 network model with intrinsic oscillation, and such network model reproduces hippocampal replay in 1-D and 2-D spatial structures. We also prove that our model functionally works as Markov-chain Monte Carlo sampling in which recall frequencies of memory patterns can be arbitrarily biased. Using this property, we implemented prioritized experience replay using our model, which actually accelerated reinforcement learning for spatial navigation. Our model explains how dynamics of hippocampal circuits realize efficient memory sampling, providing a theoretical link between dynamics and functions of hippocampal replay.

## Introduction

Hippocampus is indispensable brain region for memory formation and recall ^1^. Memory recall in the brain has been often modeled by Hopfield-type attractor network models ^2–5^. Such network models can recall memorized patterns from incomplete input patterns (auto-associative memory recall). Correspondingly, hippocampus during offline states (immobility or sleep) exhibits sequential “replay” of activity patterns experienced during tasks or mobility ^6–10^, which is necessary for learning and memory by hippocampus ^11–13^. Experiments revealed that hippocampal replay consists of sequential transitions between discrete memory patterns, which implies the existence of auto-associative memory recall in the offline hippocampus ^14,15^. Theoretically, we can implement sequential transitions between attractors by incorporating spike-frequency adaptation ^16–18^ or short-term synaptic depression ^19,20^ into attractor networks. Such network models have succeeded in reproducing replay of hippocampal place cells ^16,19,20^ and free recall of word lists ^17,18^, suggesting that attractor networks with additional instability are plausible dynamical models of memory recall in the offline hippocampus.

On the other hand, another line of research proposes functional theories of hippocampal replay from the perspective of learning efficiency. If offline hippocampal memory recall affects the value evaluation for future decision making, hippocampal replay should prioritize sampling of experiences that are useful for updating value functions. This principle corresponds to prioritized experience replay in the literature of machine learning ^21^, and studies in neuroscience have shown that the principle of prioritized experience replay explains experimental observations of hippocampal replay ^13,22–24^ (but also see ^25^). Such studies suggest that hippocampus is working as an efficient sampler of memories that supports improved learning speed and decision making.

Although these two lines of research provide bottom-up (dynamical) and top-down (functional) explanations of how hippocampal replay, the relationship between two theories is still unclear. It is unknown how we can design hippocampal attractor network models that perform adequate sampling of memories that accelerate offline learning, and how modulating the dynamical parameters of the network affects memory sampling. Bridging between hippocampal dynamics and functions not only contributes to better understanding of hippocampal mechanisms but also may lead to methods for efficient memory recall useful for various applications.

In this paper, we propose momentum Hopfield model, which is an extension of recent Hopfield-type network model ^2^ by adding momenta and kinetic energy. In contrast to conventional associative memory models that minimize the potential energy, dynamics of momentum Hopfield model conserve the sum of potential and kinetic energies. From this assumption, we derived CA3-CA1 network model with intrinsic oscillation, which is consistent with the anatomical structure ^26^ and oscillatory dynamics ^27–30^ of the hippocampus. We show that our model can reproduce sequential replay in 1-D ^6–8^ and 2-D ^9,10^ spatial structures. Furthermore, we show that we can regard sequential memory recall by momentum Hopfield model as Markov chain Monte Carlo sampling using Hamiltonian dynamics (Hamiltonian Monte Carlo sampling) ^31,32^, which has been utilized in modeling of cortical and continuous recurrent dynamics ^33,34^ and synaptic sampling ^35^. Using this property of the model, we designed and implemented biased memory sampling for reinforcement learning, which actually accelerated the learning of spatial navigation in the grid-world environment. Overall, we demonstrate that a novel theoretical framework of Hopfield dynamics with the conservation of energy theoretically links oscillatory attractor dynamics and efficient sampling of memories in hippocampus.

## Results

### Modern Hopfield model

First, we introduce the modern Hopfield model ^2^ as a conventional Hopfield-type model, which was originally proposed to interpret self-attention in transformer models ^36^ as attractor networks. Defining a *D*-dimensional state vector **x** and memory patterns **μ**_*i*_ (*i* = 1, …, *M*), the energy function of the modern Hopfield model is written as

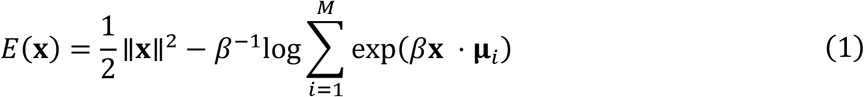

Here, inessential terms in the original definition were omitted. The original study ^2^ derived an update of **x** that resembles dot-product attention as

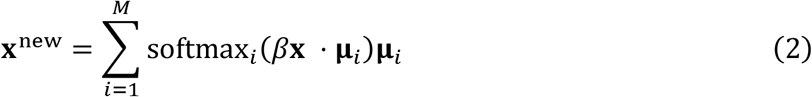

where

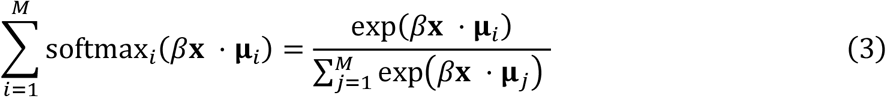

This update rule is called the modern Hopfield model because it works as an associative memory recall through attractor dynamics. Alternatively, we can apply a simple gradient descent of the energy function to derive the continuous form

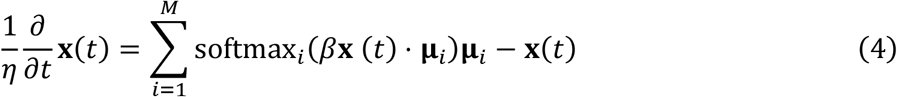

where *η* is the update rate. This update gradually changes the network state **x** such that the energy decreases.

In conventional Hopfield-type models including the modern Hopfield model, updates of the state basically minimize the energy. After several iterations, the state gets to one of minima of the energy distribution (Figure 1A, top) which correspond to embedded memory patterns (or metastable patterns in some conditions) (Figure 1A, bottom, Figure 2A). After convergence, the state becomes stable and the recalled memory is kept. This dynamical behavior is interpreted as the process of memory recall.

**Figure 1.**
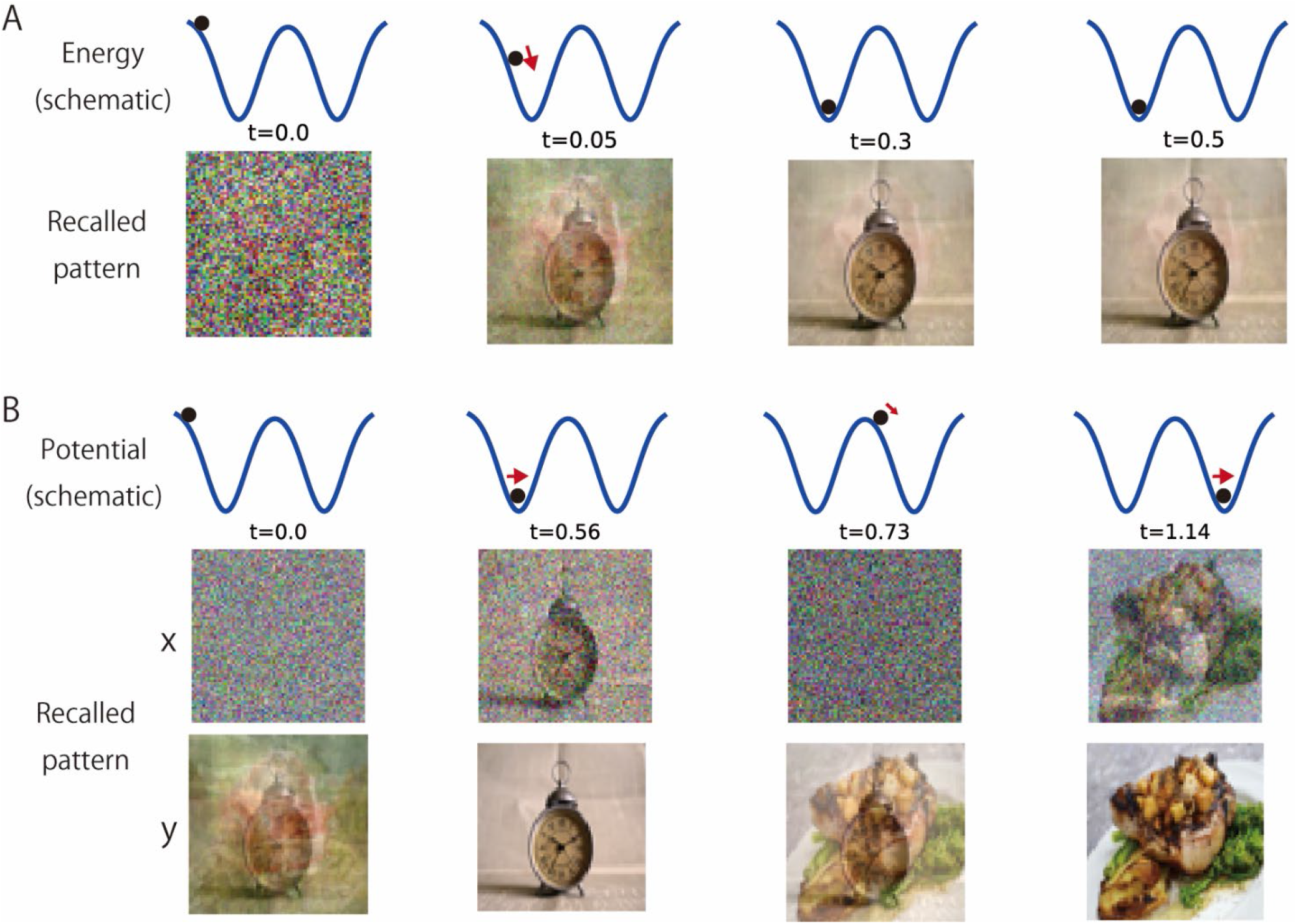
Comparison of dynamics of the modern Hopfield model and the momentum Hopfield model. Ten images (flattened and normalized pixel values) were embedded as memory patterns in each model (see Appendix 10 for the detailed methods). Black points and red arrows indicate positions and movements of state vectors, respectively. (A) A schematic of the energy landscape (top row) and recalled patterns (bottom row) in the simulation of modern Hopfield model. (B) A schematic of the potential energy landscape (top row) and recalled patterns (middle and bottom rows) in the simulation of the momentum Hopfield model.

**Figure 2.**
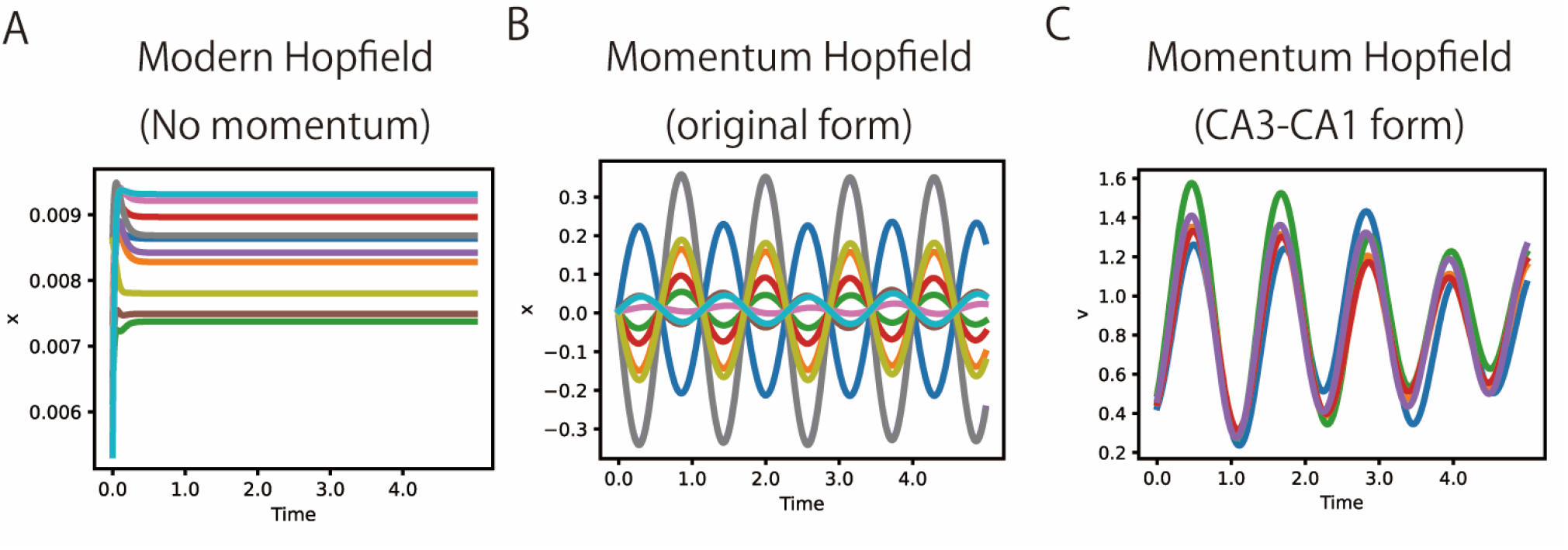
Oscillations in the momentum Hopfield model. All results correspond to the simulations in Figure 1 and Supplementary Figure 1. (A) Time evolution of 10 dimensions of **x**(*t*) in the modern Hopfield model. (B) Time evolution of 10 dimensions of **x**(*t*) in the momentum Hopfield model (original form). (C) Time evolution of *v*_*k*_(*t*) in the momentum Hopfield model (CA3-CA1 form).

### Momentum Hopfield model

We propose the momentum Hopfield model by extending the modern Hopfield model with Hamiltonian dynamics. Here, we regard the state vector **x** and the energy function *E*(**x**) in the modern Hopfield model as a generalized position and potential energy, respectively. We add generalized momentum vector **r** and kinetic energy *K*(**r**) to define the following Hamiltonian (the total energy)

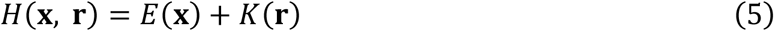

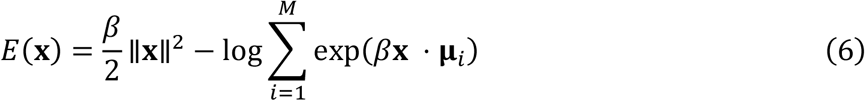

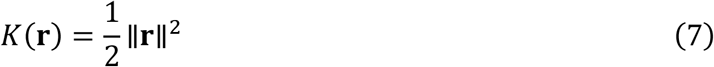

The momentum Hopfield model is derived as canonical equations for the Hamiltonian:

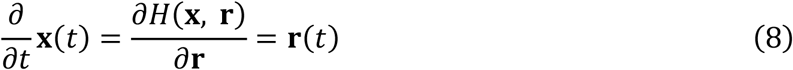

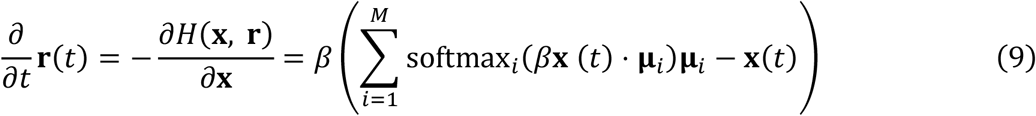

Hamiltonian *H*(**x, r**) is kept constant in the temporal evolution of (**x, r**) following these differential equations that conserves the total energy. The conservation of the total energy was also guaranteed in numerical simulations in this paper by using leap-frog integration ^32^ (see Methods).

In contrast to the conventional Hopfield-type models, temporal evolution of (**x, r**) in the momentum Hopfield model conserves the sum of potential and kinetic energies. Therefore, as the state falls into a well of the potential energy to recall a memory, kinetic energy increases and makes the state escape from the potential well and travel to another close potential well (Figure 1B, top). Such dynamics result in transitions across multiple similar memory patterns, which can be interpreted as sequential memory recall. We note that this model no longer has fixed-point attractors in a strict sense, similarly to attractor networks with additional instability such as adaptation and short-term depression ^16–20^.

However, **x** in the momentum Hopfield model is often very noisy (Figure 1B, middle). Therefore, we introduce an additional readout vector **y** for precise memory recall as

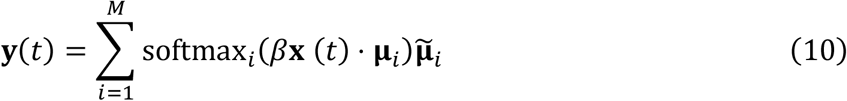

This readout does not affect the internal dynamics of memory recall but enables precise recall of embedded memory patterns (Figure 1B, bottom; Supplementary video 1). Here, we additionally introduced vectors 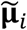 to enable arbitrary hetero-associative memory recall (key-value memory) by the model ^2,3,36^. Although these readout patterns can be arbitrarily defined, we basically assume 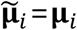 which just projects **x**(*t*) to close patterns unless otherwise specified.

By eliminating **r**(*t*), the momentum Hopfield model can be simplified as

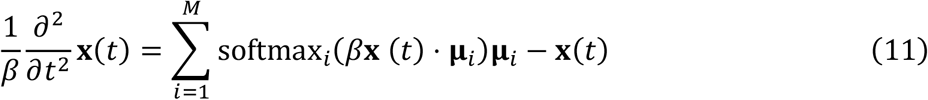

Therefore, the momentum Hopfield model is the extension of the modern Hopfield model (in the continuous form) by substituting the first-order time derivative 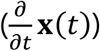 by the second-order time derivative 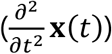. It is notable that this second-order dynamics similar to a spring-mass system 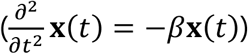 generate intrinsic oscillation in the model (Figure 2B). Our model suggests that such oscillation can facilitate sequential memory recall of Hopfield-type model.

### Relating the model to the structure of hippocampal circuits

In this section, we argue the anatomical relevance of the proposed model with hippocampal circuits. The modern Hopfield model can be interpreted as two-layer neural network in which **x**(*t*) are feature neurons and softmax layers as memory neurons ^5^. Although this interpretation is applicable to our model (Figure 3, left), we provide a more specific correspondence between the model and the hippocampal structure.

**Figure 3.**
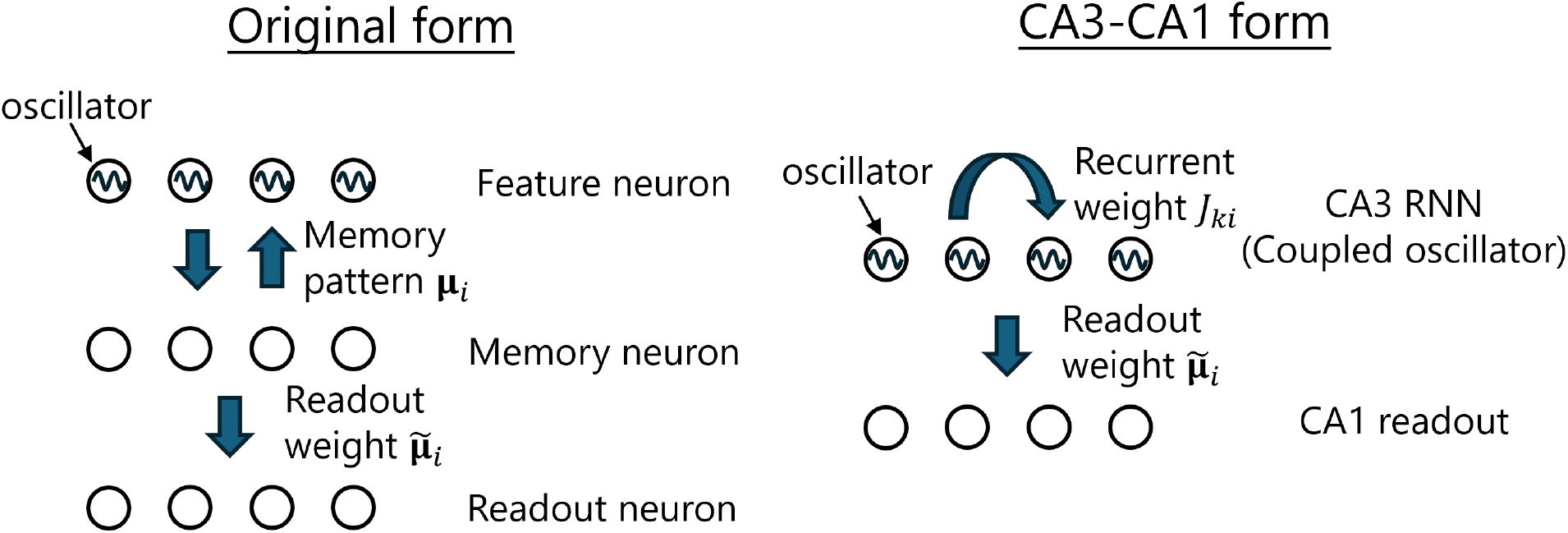
Network structures of momentum Hopfield model in the original form (left) and the CA3-CA1 form (right).

Here, we focus on two major hippocampal subfields, CA3 and CA1. CA3 has plenty of recurrent synapses, and CA1 receives many feed-forward synaptic inputs from CA3 (Schaffer collaterals) ^26^. Previous studies suggest that CA3 recurrent networks provide attractor dynamics for memory recall and generate sequential replay ^37–40^, and CA1 acts as readout of CA3 activities.

By multiplying **μ**_*k*_ (*k* = 1, …, *M*) to the momentum Hopfield model, we can obtain the recurrent network model with readout as follows:

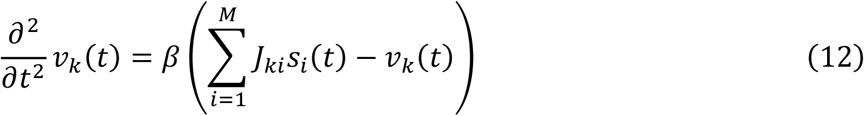

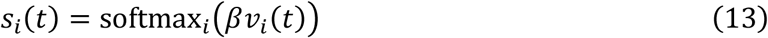

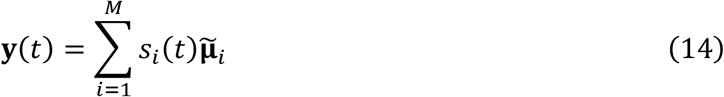

where *v*_*k*_(*t*) = **x**(*t*) ⋅ **μ**_*k*_ and *J*_*ki*_ = **μ**_*k*_ ⋅ **μ**_*i*_. Here we can regard *v*_*k*_(*t*), *s*_*i*_(*t*), *J*_*ki*_ as the internal activation (membrane potential), the neural activity (firing rate), and recurrent synaptic weights of CA3 neurons. Then, softmax_*i*_(*βv*_*i*_(*t*)) corresponds to the nonlinear rate function that is normalized by divisive inhibition. Also, **y**(*t*) and 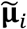 correspond to CA1 neural activities and synaptic weights of Schaffer collaterals. In this form, we can relate the momentum Hopfield model to the anatomical structure of CA3-CA1 network (Figure 3, right).

We call the equations before the transformation as the original form (Figure 3, left), and the equations after transformation as the CA3-CA1 form (Figure 3, right). The CA3-CA1 form can perform memory retrieval similarly to the original form (Supplementary Figure 1), and the internal activation in each CA3 neuron (*v*_*k*_(*t*)) exhibits an intrinsic oscillation (Figure 2C). Therefore, the CA3 network can be interpreted as a coupled oscillator in this model (Figure 3, right). This oscillation possibly corresponds to gamma oscillation in CA3 synchronized with sharp-wave ripple events ^27–30^ (but see Discussion for several other interpretations). We use the CA3-CA1 form in the following simulations unless the analysis of **x**(*t*) is necessary.

### Replay of place cells in 1-D spatial structures

The sequential memory recall by the momentum Hopfield model is reminiscent of sequential replay of place cells in hippocampus. Therefore, we checked whether momentum Hopfield model can reproduce hippocampal replay of 1-D and 2-D spatial structures.

First, we simulated replay of place cells on a linear track ^6–8^. Gaussian place fields are uniformly distributed on the track (Figure 4A), and we created memory patterns **μ**_*i*_ and recurrent weights *J*_*ki*_ by sampling population activity patterns of place cells at various locations on the track (Figure 4B and 4C) (see Methods for the detail). We simulated the momentum Hopfield model in this setting and performed Bayesian place decoding ^14,15^ of **y**(*t*) to visualize how place information is recalled in the simulation (see Methods for the detail). Then, we could produce sequential replay of place cells corresponding to the run on the linear track (Figure 4D and 4E). When the number of memories is large (8 memory patterns), replay is apparently continuous (Figure 4D). In contrast, when the number of memories is small (4 memory patterns), replay exhibits jumps across discrete memory patterns (Figure 4E) which is reminiscent of experimental observations in the hippocampus ^14,15^. As in the simulation of image retrieval, the internal activation of CA3 neurons (*v*_*k*_(*t*)) exhibits oscillations during the generation of sequential replay (Supplementary Figure 2). This result demonstrates the emergence of sequential replay of place cells by the momentum Hopfield model.

**Figure 4.**
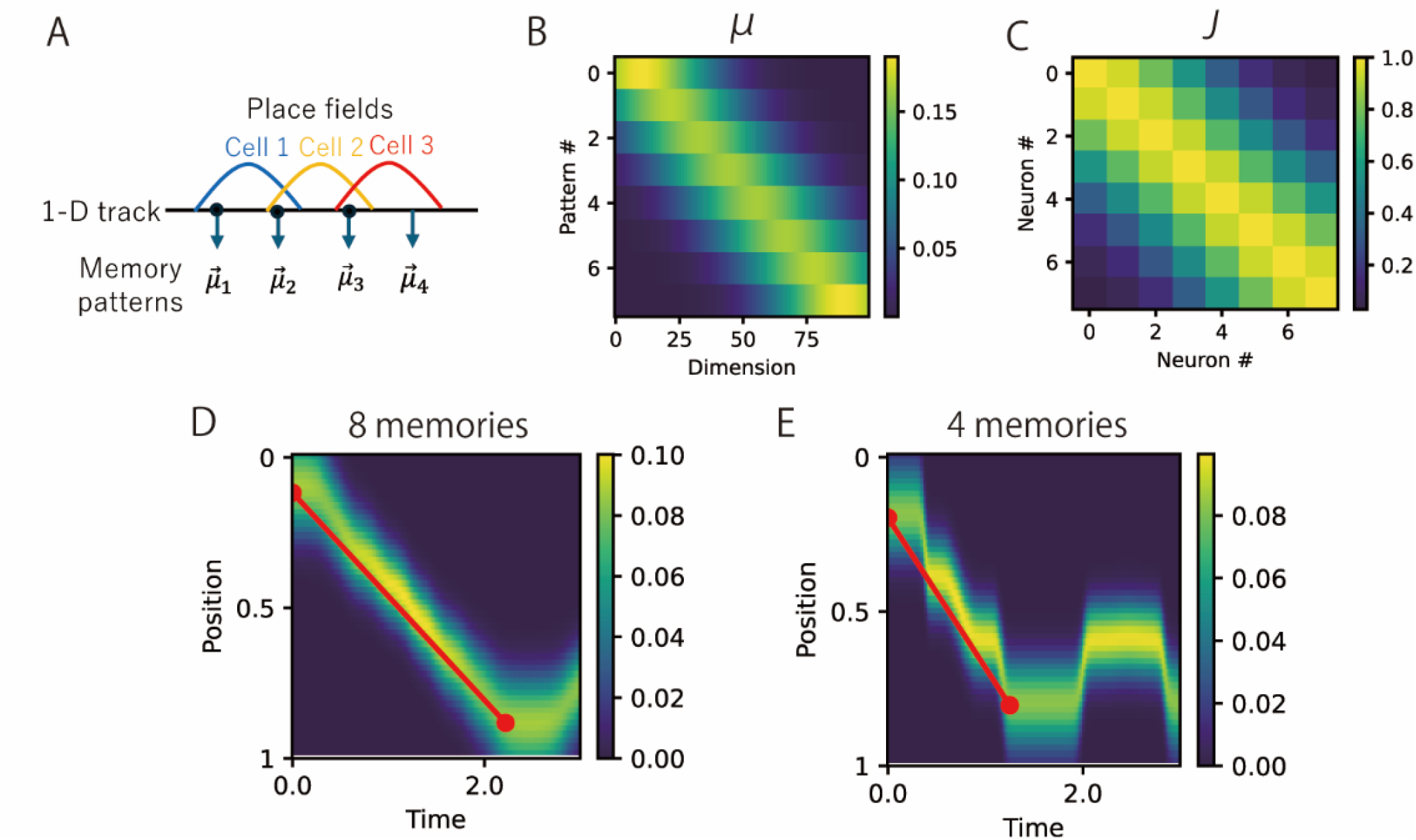
Simulations of replay of place cells on a linear track. (A) A schematic of the simulation setting. 100 neurons have place fields uniformly distributed on the linear track. Memory patterns were created from population activity patterns of place cells at several locations on the track. (B) Memory patterns used in the simulation (8 memory patterns). (C) Recurrent weight matrix **J** in the CA3-CA1 form. (D) Sequential replay in the momentum Hopfield model with 8 memory patterns. Heatmap shows probabilities of positions at each time that were obtained by Bayesian place decoding of simulated activity patterns. A red line indicates a linear fit between the start and the end of replay. (E) Sequential replay in the momentum Hopfield model with 4 memory patterns.

### Replay of place cells in 2-D spatial structures

Next, we simulated replay of place cells in a 2-D space (corresponding to a square chamber) ^9,10^. As in the linear track, we assumed place fields are uniformly distributed in the 2-D field (Figure 5A), and sampled memory patterns for momentum Hopfield model at various locations in the field (see Methods for the detail). Then, we performed Bayesian place decoding of simulated activity patterns of the model, and visualized replay trajectories (trajectories of the gravity center of the decoded probability map at each time). Then, we could see that the model generated sequential replay in the 2-D space (Figure 5B, Supplementary video 2).

**Figure 5.**
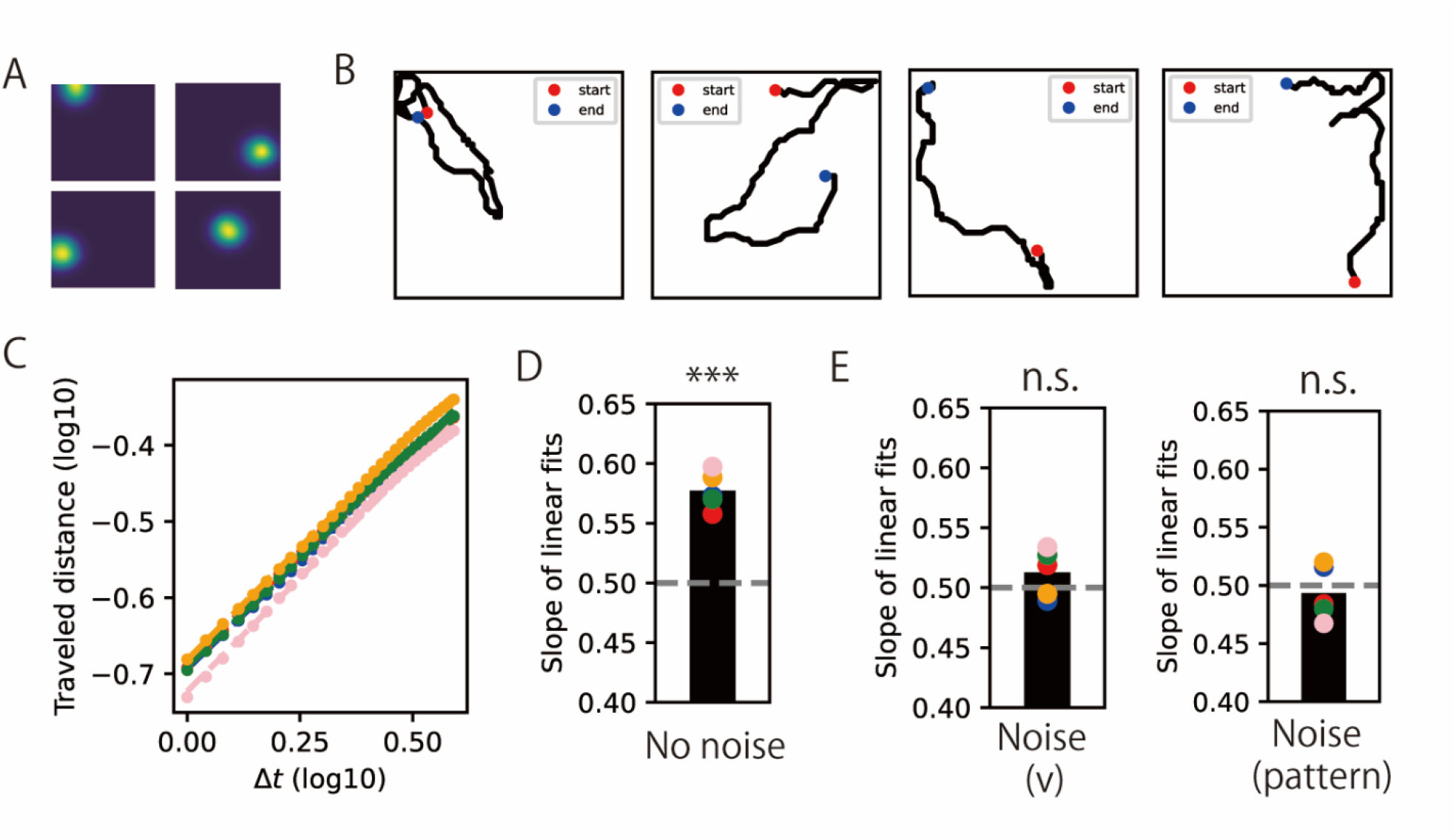
Simulations of replay of place cells in a 2-D field. (A) Example place fields in the simulation. (B) Example replay trajectories (the gravity center of probability maps of Baysiean place decoding at each time) generated by the momentum Hopfield model. (C) Log-log plot of the relationship of time difference and traveled distance in replay trajectories. Each color indicates an independent simulation using a different random seed. (D) Slopes of linear fits of data in (C). Points indicate independent simulations and the bar indicates the mean of five simulations. The slope was significantly different from 0.5 (p<0.001; five samples). (E) Slopes of linear fits of log-log plot in the simulations with noise to **v**(*t*) (left) and **μ**_*i*_ (right). In both cases, slopes were not significantly different from 0.5 (five samples). All statistical tests were two-sided one-sample t-test with Bonferroni correction.

Although such sequential replay in 2-D spaces is observed in the rat hippocampus, replay trajectories exhibit different properties between sleep and awake periods. During sleep, replay trajectories have a statistic property of Brownian motion, that is, time differences (*t*) and traveled distances (*d*(*t*)) satisfies the relationship 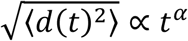 where *α* = 0.5 ^9^. In contrast, replay trajectories during awake periods can be fitted well by the model with momentum in the space, and do not satisfy a property of Brownian motion (*α* > 0.5) ^10^. Because our model is driven by momentum in the representation space, we can expect that replay trajectories in our model have the property during awake periods. Consistent with this expectation, time differences and traveled distances in the simulation show linear relationships in log-log plot (Figure 5C) and the slope (*α*) was significantly larger than 0.5 (Figure 5D). This is the result of the deterministic simulation, but adding noise in the model may lead to Brownian-motion-like replay trajectories during sleep. We tested adding noise to either the internal activation *v*_*k*_(*t*) or the pattern **μ**_*i*_, and confirmed the emergence of Brownian-motion-like replay (*α* ≈ 0.5) in both cases (Figure 5E). These results suggest that the difference of replay trajectories between awake and sleep states can be explained by noise level in the hippocampus.

In summary, by embedding activity patterns of place cells in 1-D and 2-D spatial structures, the momentum Hopfield model produces sequential 1-D replay and 2-D replay which are experimentally observed in hippocampus.

### Momentum Hopfield model performs Markov chain Monte Carlo sampling

So far, we have seen the biological relevance of the momentum Hopfield model to hippocampal replay. Next, we see a functional implication of the model. Specifically, we show that the momentum Hopfield model performs Hamiltonian Monte Carlo sampling (HMCS) (or Hybrid Monte Carlo sampling) ^31,32^, one of efficient methods for Markov chain Monte Carlo sampling. From this theory, we can rigorously calculate how memory sampling occurs in the momentum Hopfield model.

HMCS utilizes Hamiltonian dynamics with potential energy *E*(**x**) to perform Markov-chain Monte Carlo sampling from probabilistic distributions 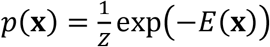 (*Z* is a normalizing constant). Details of HMCS are described in Methods. We consider HMCS that samples from the following Gaussian mixture in the N-dimensional space

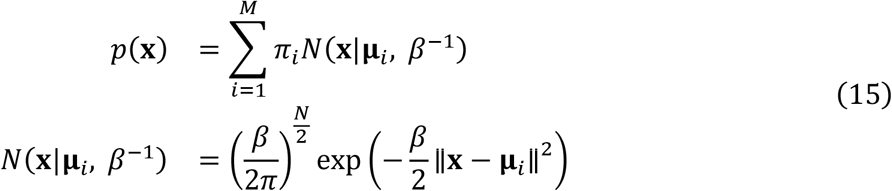

where i-th Gaussian distribution has the mean parameter **μ**_*i*_ (*i* = 1, …, *M*), and isotropic variance *β*^−1^, and the mixing ratio *π*_*i*_.This distribution can be transformed into the form 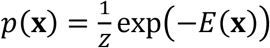 by setting potential energy as

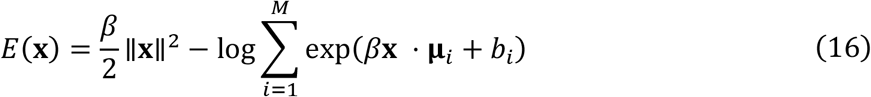

where 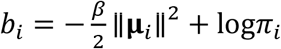. By setting 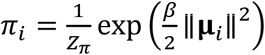 (*Z*_*π*_ is a normalizing constant), *b*_*i*_ = −log*Z*_*π*_ becomes constant and is absorbed into *Z*. Then, *b*_*i*_ disappears from the energy and we can obtain exactly same potential energy as the momentum Hopfield model. This suggests that sequential memory recall by the momentum Hopfield model can be interpreted as sampling from a Gaussian mixture in which means **μ**_*i*_ correspond to embedded memory patterns. If bias parameters *b*_*i*_ is zero, expected frequencies of memory recall is solely determined by norms of embedded patterns as 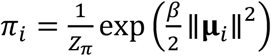 which predicts frequencies of memory recall by the model. Conversely, we can arbitrarily control frequencies of memory recall by introducing additional bias parameters 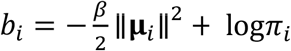.

To check that the momentum Hopfield model actually performs sampling from Gaussian mixtures, we performed simulations in 2-D state space (**x**(*t*) ∈ ℝ^2^; note that this is not a physical 2-D space and we did not use place cells) (Figure 6A and 6D). In the simulation, we varied the mixing ratio of Gaussian mixture by two different ways. First, L2 norm of all memory patterns ∥**μ**_*i*_ ∥^2^ are normalized and bias parameters are modulated as 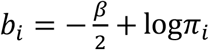 (Figure 4A-C). Second, bias parameters are removed (*b*_*i*_ = 0) and L2 norm of embedded memory patterns ∥**μ**_*i*_∥^2^ (Figure 6D-F) of the momentum Hopfield model. In both cases, activity patterns of the momentum Hopfield model approximates theoretically expected Gaussian mixture distributions (Figure 6B and 6E) and the recall frequency matched the theoretical prediction (Figure 6C and 6F).

**Figure 6.**
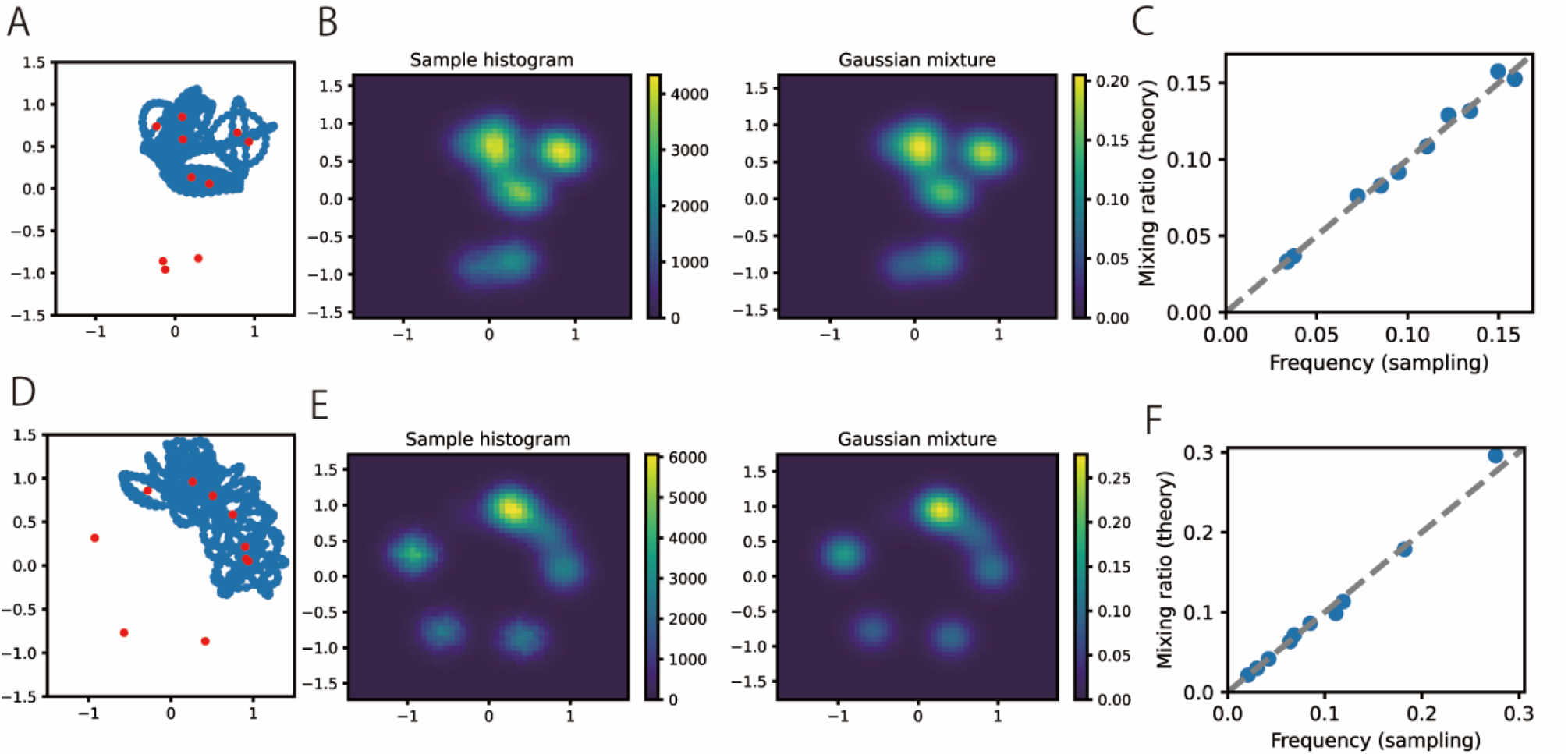
Sampling of Gaussian mixture in a 2-D state space by Momentum Hopfield model. We note that the plotted space is not a physical 2-D space but the space of 2-D state vector (***x***(*t*) ∈ ℝ^2^). (A,B,C) A simulation in which bias parameters were modulated. (A) Embedded memory patterns (red points) and a part of the simulated trajectory of the state vector ***x***(*t*) (a blue line). (B) Histogram of sampled state vector ***x***(*t*) (left) and the theoretically expected distribution (Gaussian mixture) (right). (C) Relationship between theoretical and empirical sampling frequency of each memory pattern. (D,E,F) A simulation in which L2 norms of embedded memory patterns were modulated.

We also checked that the theory of sampling discussed above can be used to create biased replay of place cells in the 2-D field. We modulated L2 norms of a part of memory patterns in the simulation in the square 2-D space (as in the previous section) and confirmed that the sampling frequency of memory patterns are biased as theoretically expected (Figure 7).

**Figure 7.**
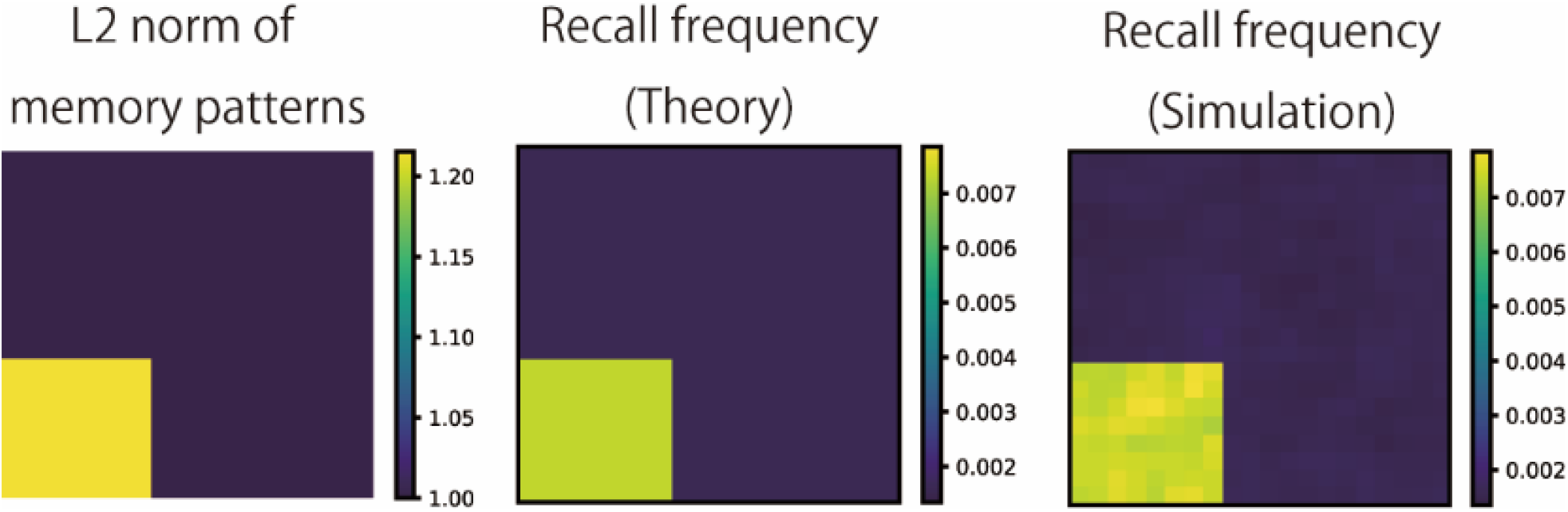
Biased recall frequencies in replay of 2-D place cells. By changing L2 norms of memory patterns in a specific region (left), theoretical expectation of recall frequency is modulated (middle). Recall frequency in the simulation follows the theory (right).

### Generating prioritized experience replay for reinforcement learning

Finally, we implemented memory sampling using the momentum Hopfield model that accelerates value learning for spatial navigation as prioritized experience replay ^13,21–24^. Here we consider the momentum Hopfield model in which sampling frequency follow 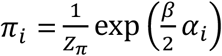, and modulate the tunable parameter *α*_*i*_ by various factors such as TD errors. Such prioritization is possible through modulation of L2 norms of memory patterns as 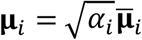 where 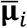 is unit-norm pattern vectors, and this setting leads to the network model with amplification of inputs and outputs of each CA3 neuron by 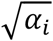 (see Methods). This modulation can be biologically implemented as dopaminergic amplification of hippocampal activities ^41–44^ (see Discussion for other possibilities).

To check whether replay in such model accelerates learning, we evaluated the performance of reinforcement learning of spatial navigation in a 2-D grid-world (Figure 8A). We performed Q learning of value functions through sampling (replay) of episodes in the grid-world. Each episode is a set of a current position (state) *s*, a next position *s*_next_, a movement to one of four directions (action) *a*, and reward *r*. To perform sampling, we embedded memory patterns corresponding to episodes in the momentum Hopfield model. We created memory patterns using disentangled successor information (DSI) ^45^ which generates hippocampus-like representations based on successor representation (SR) ^46,47^. This model represents episodes that are spatially close (in other words, those often visited within a short time window) by similar representation vectors. Although they are representations of episodes (state-action pairs) but not spatial positions, we could see spatial representations like place cells by taking the average with respect to actions (Figure 8B). Furthermore, the momentum Hopfield model with those memory patterns produced sequential replay in the 2-D grid-world (Figure 8C). See Methods for the detail of reinforcement learning and sampling of episodes by the momentum Hopfield model.

**Figure 8.**
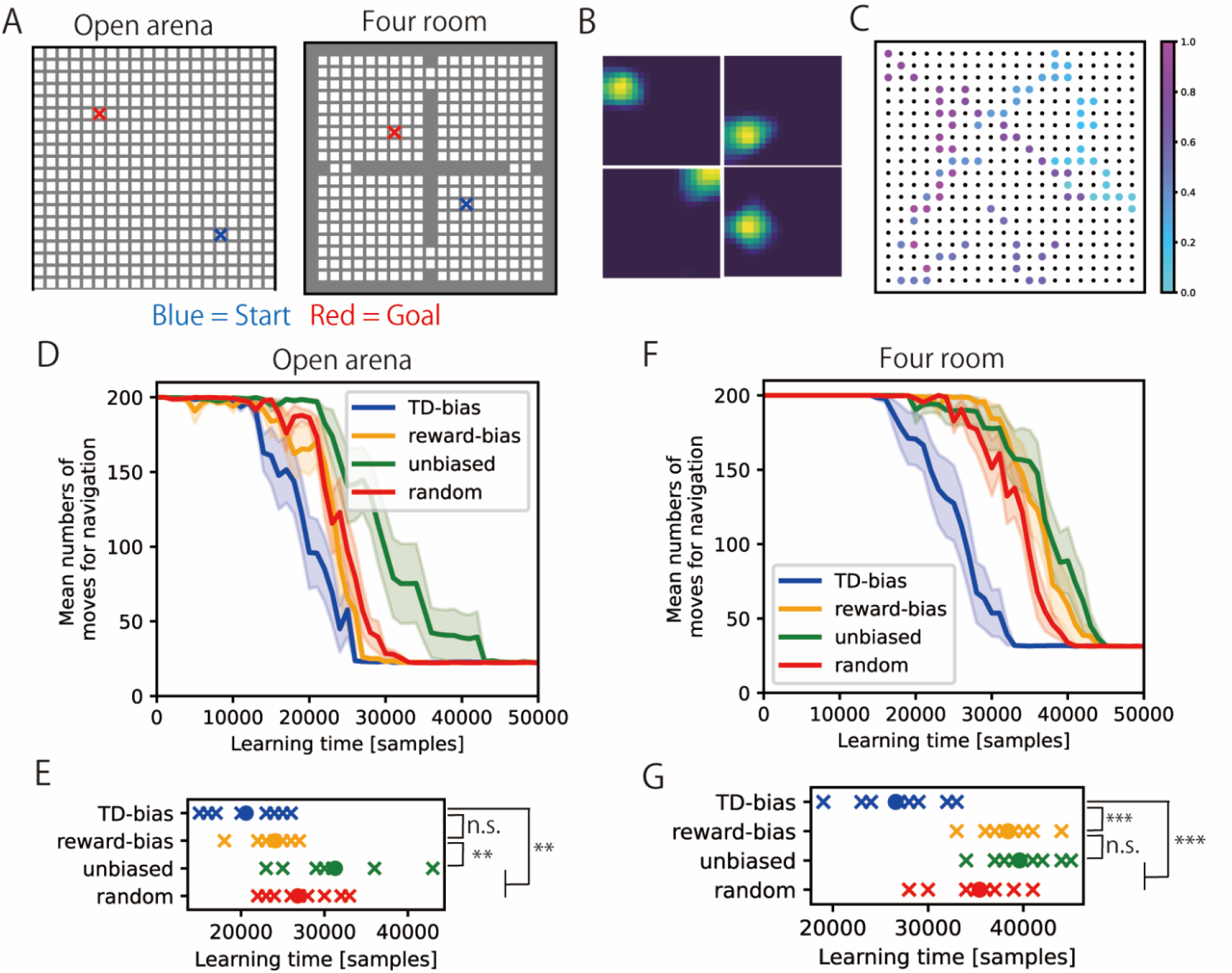
Prioritized experience replay by the momentum Hopfield model. (A) The setting of the spatial navigation task in a 20 x 20 grid world. Opena arena (left) and four separate rooms (right). (B) Examples of spatial representations in memory patterns created by DSI-sparse. We took mean activities of episodes that has the same *s* (current position). (C) An example of replay trajectory during learning (TD-bias). We plotted positions corresponding to *s* (current position) in sampled episodes. Black dots indicate 20 × 20 grid world and the color indicate time in the plotted segment. (D) The performance of spatial navigation in the open-arena setting (the number of moves from start to goal). The performance was evaluated at different times during learning. Mean (dark-colored lines) and standard deviation (pale-colored area) for each setting were calculated from ten independent simulations with different random seeds. (E) Comparison of learning time (the number of samples when the number of moves fell below 40) by four replay methods in the open-arena setting. We performed 10 independent simulations in each condition. (F) The performance of spatial navigation in the four-room setting (same as D). (G) Comparison of learning time in the four-room setting. All comparisons were two-sided two-sample t-test with Bonferroni correction; ** p<0.01, *** p<0.001.

Here, we tested two strategies for modulation of the parameter *α*_*i*_. First, as in the original prioritized experience replay in RL ^21^, the importance of each memory is determined by the absolute TD error in the past TD learning (TD-bias). Second, as in the previous biologically plausible implementation of prioritized replay ^24^, the importance of each memory is determined by the proximity to reward (reward-bias). We compared the performance of these two methods with the momentum Hopfield model without modulations of recall frequencies (unbiased) and uniformly random sampling of experiences without the momentum Hopfield model (random).

As the result, TD-bias accelerated the convergence of value functions (Supplementary Figure 3) and the improved the learning speed of spatial navigation (Figure 8D, 8E, 8F, 8G) than unbiased and random sampling. Reward-bias setting accelerated learning in the relatively easy open-arena setting (Figure 8D, 8E) whereas reward-bias did not improve the learning in the four-room setting (Figure 8F, 8G). This implies that biasing the sampling by proximity to reward only improves learning in easy settings in which the shortest path length is small.

We can understand this difference between TD-bias and reward-bias by analyzing changes of sampling frequencies during learning. As expected, reward-bias always selectively samples around reward positions (Supplementary Figure 4). Although this strategy accelerates updates of value functions at the beginning, learning becomes slow at the end of learning when updates at distant positions from reward is necessary. In contrast, TD-bias selectively replays positions around reward at the beginning and then changes the strategy to unbiased and uniform sampling of the whole space (Supplementary Figure 4). Such flexible replay strategy enables fast value updates both at the beginning and the end of reinforcement learning.

In summary, we can implement efficient memory sampling for reinforcement learning by appropriate modulation of replay frequency in momentum Hopfield model, which provides a way to implement hypothesized functions of hippocampal replay, prioritized experience repaly, by extended attractor dynamics.

## Discussion

In this paper, we proposed the momentum Hopfield model, which extends conventional Hopfield-type attractor networks with momentum, kinetic energy, and conservation of total energy. The model can be interpreted as a CA3-CA1 network with intrinsic oscillation, and the model successfully reproduced sequential replay for 1-D and 2-D spatial structures. Furthermore, we provided a functional interpretation of our model as a Monte Carlo sampler. Using this theory, we demonstrated the implementation of prioritized experience replay by hippocampal network model. Overall, our model connects two lines of theoretical research about dynamical (Hopfield-type attractor dynamics and oscillation) and functional (efficient memory sampling for reinforcement learning) aspects of hippocampus.

### Biological interpretations of the momentum Hopfield model

Through the transformation of the momentum Hopfield model, we provided the interpretation of the model as the CA3-CA1 circuit in the hippocampus. The internal dynamics in CA3 neurons (*v*_*k*_(*t*)) are given by the second-order differential equation. This biological interpretation necessitates oscillatory dynamics of CA3 neural units (neurons or neural subpopulations in CA3), and the CA3 network should work as a coupled oscillator network. One of biologically plausible interpretations of this oscillation is slow gamma oscillation that coincides with sharp-wave ripple events ^27–30^. Slow gamma oscillation originates in CA3 ^27,28^ and deeply relates to replay dynamics in 1-D and 2-D spatial structures ^9,14,15^ (but there is controversy on the existence of slow gamma oscillations ^48^). The source of gamma oscillation can be an interaction between excitatory and inhibitory neural populations ^49–51^, which are implementable in local circuits of CA3. Otherwise, oscillation can be generated by intracellular mechanisms ^52^. Furthermore, it is also possible to relate our model to theta and ripple-band oscillations in the hippocampus ^27,40,53^. Our model currently does not exclude these multiple possibilities because the current model lacks details of biological neurons and synapses. Implementation of our model with detailed neuron models and validation with the real neural data are left for future studies.

In the simulation of prioritized experience replay, we demonstrated amplification of inputs and outputs of CA3 neurons by absolute TD errors leads to accelerated value learning. This amplification may correspond to dopaminergic modulations in the hippocampus. However, reward-related dopaminergic effects have been mostly studied in CA1 ^41–44^ whereas CA3 receives a dopaminergic innervation from locus coeruleus which signals novelty ^54^. Therefore, biasing replay in CA3 may be affected by novelty rather than reward-related information. Such novelty-dependent modulation is consistent with the increase of replay during sleep after novel experiences ^55–57^, which can support efficient memory consolidation. However, other studies suggest that dopaminergic innervation to dentate gyrus relates to reward information and reinforcement learning ^58–60^. Therefore, it is possible that reward-related amplification in dentate gyrus affects CA3 dynamics through modulation of bias parameters *b*_*i*_ rather than norms of memory patterns **μ**_*i*_. Such bias parameters may be useful to build more realistic hippocampal models although such parameters were not considered in the original modern Hopfield model ^2^.

In our simulations, we demonstrated that our model without noise generated non-Brownian-motion replay dynamics observed in the awake state ^10^, and adding noise yielded Brownian-motion-like replay dynamics observed in sleep ^9^. These results suggest that there should be difference in noise level between awake and sleep states. It is natural that replay during sleep is driven by noise because of lack of sensory inputs, and consistently, replay fidelity is lower during sleep than during awake ^61^. Furthermore, we may also interpret this difference from the functional point of view. The role of awake replay should be task-specific and reflect the behavioral trajectories of animals which are not Brownian motion ^9,10^, whereas replay during sleep presumably contributes to memory consolidation ^11,62^ for which random activation is useful enough.

### Theoretical implications of the momentum Hopfield model

In this paper, we gave an interpretation of the momentum Hopfield model as a sampler using the theory of Hamiltonian Monte Carlo sampling (HMCS). HMCS has been used to model dynamics in primary visual cortex ^34^, continuous (bump) attractor networks ^33^, and recurrent networks for path integration ^63^. This line of research consistently proposes that dynamic activity patterns in the brain are the result of sampling. Sampling is done by second-order Hamiltonian dynamics, which enables more efficient sampling than sampling based on random walk ^31,32^. Our model applies this view to hippocampal replay, and proposes that hippocampal dynamics acts as an efficient memory sampler. Although there are various interpretations of momentum variables as activities of interneurons ^34^ and adaptation ^33,63^, it is unclear whether such interpretations are applicable to our model based on Hopfield-type dynamics.

A concurrent work from other authors demonstrated that path integration networks with adaptation generates efficient hippocampal replay with momentum ^63^. Similarly to our model, they propose that hippocampal replay may be driven by second-order dynamics, and such dynamics support efficient memory sampling. However, their model is obtained through training of normal recurrent networks for path integration in contrast that our model is based on the combination of the theory of Hopfield-type attractor networks with oscillation. These studies suggest that both oscillatory auto-associative dynamics in CA3 and path integration in the entorhinal cortex^64,65^ can support replay dynamics with momentum.

Based on the theoretical interpretation of the momentum Hopfield model as a sampler, we designed the neural network model for prioritized experience replay. However, our theory can be generally applied to biological implementation of other probabilistic computations. For example, we may use our model to implement Bayesian inference like previous Hamiltonian-based models ^33,34^. Specifically, hippocampus is often related to sequential Bayesian inference and hidden Markov models (HMM) ^66^. Our theory may provide a way to implement such HMM-based hippocampal models using oscillation.

We derived the momentum Hopfield model from modern Hopfield model ^2^, which has the same mathematical form with a self-attention module in transformer-type models ^36^. Therefore, it is mathematically straightforward to relate our model to self-attention. Previous studies have shown the relationship between self-attention and hippocampal memory recall: some attention heads in transformer-based large language models mechanistically resemble human episodic memory ^67^, and self-attention can exhibit spatial representations like hippocampus in spatial learning ^68^. However, sequential replay observed in hippocampus has not been related to the mechanics of self-attention and it is unclear whether there are merits to implement sequential replay in such large-scale predictive models. This problem is intriguing to be explored in the future.

## Methods

### Simulation of the momentum Hopfield model

To perform numerical simulations, we derive a discretized form of the model by leap-frog integration ^32^ which guarantees the conservation of Hamiltonian through the temporal evolution. With a step size *ϵ*, leap-frog integration of **x** and **r** in the original form is

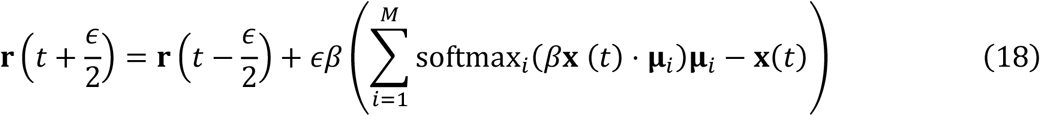

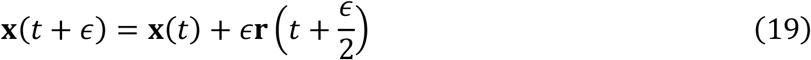

Therefore, we calculated discretized dynamics in *n* = 1,2,3, … as

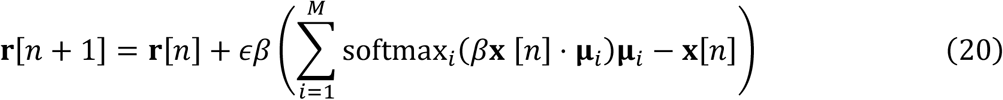

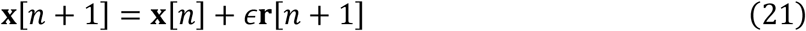

and obtained **x**(*nϵ*) = **x**[*n*] (and implicitly, **r**[*n*] corresponds to **r**(*n* − 0.5)*ϵ*).

Similarly, the simulation of the CA3-CA1 form is performed by

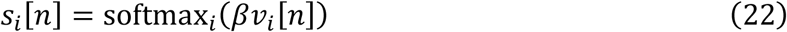

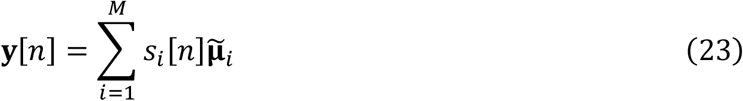

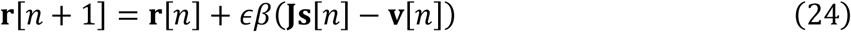

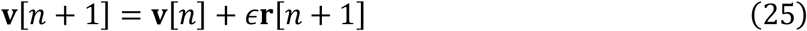

where **v** and **J** are vectors of *v*_*i*_(*t*) and *s*_*i*_(*t*), and **J** is the matrix of *J*_*ki*_. Initialization was often performed for **x**[0], then transformed to **v**[0] = **Mx**[0] where **M** = (**μ**_0_, **μ**_1_, …, **μ**_*M*_)^T^. Also, **r**[*n*] was sometimes resampled from Gaussian distribution following the requirements of Hamiltonian Monte Carlo sampling (see the explanation below).

### Hamiltonian Monte Carlo sampling

Here we briefly explain a sampling framework called Hamiltonian Monte Carlo sampling (HMCS) ^31,32^ (also called Hybrid Monte Carlo sampling).

Our goal is sampling from a probabilistic distribution *p*(**x**) using HMCS. Then, we consider potential energy *E*(**x**) that satisfies 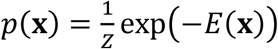 (*Z* is an arbitrary normalizing constant). We also consider kinetic energy

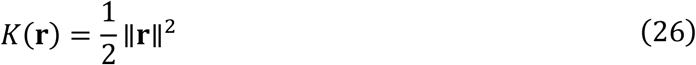

which corresponds to the Gaussian distribution

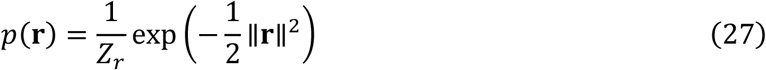

Then, we can define Hamiltonian that corresponds to the simultaneous distribution of (**x, r**)

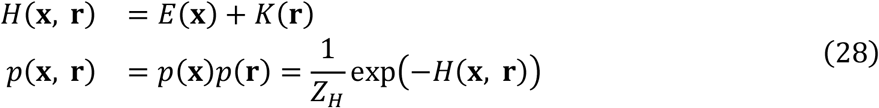

Because **x** and **r** are independent in this distribution, we can sample from *p*(**x**) by sampling (**x, r**) from *p*(**x, r**) and discarding **r**. In HMCS, (**x, r**) is sampled by numerically solving Hamiltonian dynamics

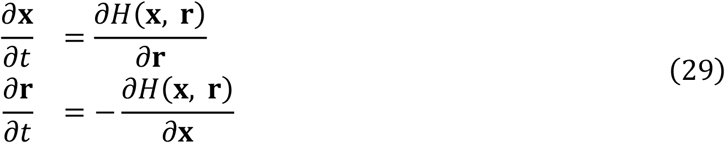

by leap-frog integration. In each update from (**x**_1_, **r**_1_) to (**x**_2_, **r**_2_), Hamiltonian is conserved (*H*(**x**_1_, **r**_1_) = *H*(**x**_2_, **r**_2_)), so *p*(**x**_1_, **r**_1_) = *p*(**x**_2_, **r**_2_) always holds (if there is no numerical error). Therefore, Metropolis acceptance criterion 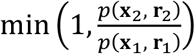 is always 1, and we can interpret Hamiltonian dynamics as Markov-chain Monte Carlo sampling for *p*(**x, r**) that is always accepted. Therefore, iteration of simulating Hamiltonian dynamics realizes efficient sampling without sample rejection. Furthermore, adding momenta and kinetic energy enables large jumps in sampling that is hardly obtained by the random walk. This property often supports efficient sampling from the whole space by small samples.

However, these updates do not realize sampling from the whole space of (**x, r**) (non-Ergodic) because *p*(**x, r**) is always kept constant. To perform Ergodic sampling, **r** is replaced by samples from Gaussian distribution *p*(**r**) after a certain length of leap-frog iteration. This operation can be regarded as Gibbs sampling, which is also Markov-chain Monte Carlo sampling with the acceptance rate 1. Such resampling is performed in most of simulations in this paper by resetting **r**[*n*] (and also **x**[*n*] and **v**[*n*] in some simulations) intermittently. Such resetting can naturally occur in the hippocampal replay because they are often intermittent bursting events synchronized to sharp-wave ripples.

Furthermore, the original HMCS has various heuristics to remove biases in sampling such as rejection of samples to remove the effect of numerical errors. However, we omitted those additional procedures in this work, which did not significantly degrade the performance at least in simulations in this paper.

### Bayesian place decoding

In simulations of place-cell replay, we performed Bayesian place decoding of simulated neural activity patterns as in previous experimental studies ^14,15^.

First, assuming that the maximum firing rate of place cells is 10 Hz and the size of time bins is 0.02 s, we transformed population activity patterns for binned positions (spatial bins) **ν**_*k*_ to firing rates **F**_*k*_ as following

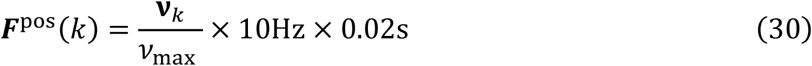

where *v*_max_ is the largest element in all **ν**_*k*_. We also performed the same transformation to simulated vectors **y**(*t*) to obtain simulated firing rates ***F***^sim^(*k*).

Next, assuming that neurons generate spikes from independent Poisson processes, and the prior distribution for positions is uniform, we calculated a posterior probability of *k*-th spatial bin at time *t P*_*k*_(*t*) as

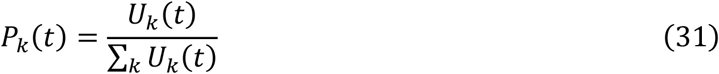

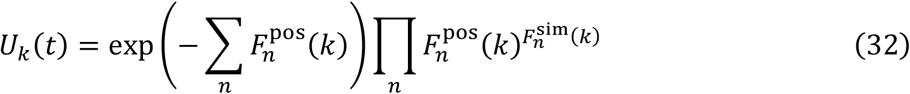

where *F_n_*(*k*) is the *n*-th element of **F**(*k*).

### Simulation methods of the example of image retrieval (Figure 1 and 2)

We used ten images from pxhere (https://pxhere.com) for the simulation. All images were resized to 64 × 64, and we flattened pixel values of each image to obtain memory pattern vectors (**μ**_*i*_). Then, memory patterns were normalized such that ∥**μ**_*i*_∥^2^ = 1. In the simulation of the modern Hopfield model, we simulated the continuous version (Eq. (4)) by Euler method (*β* = 30, *η* = 1 and the step size 0.01). In the simulation of the momentum Hopfield model, we solved leap-frog integration in the original form (Eq. (20-21)) with *β* = 30 and *ϵ* = 0.01. In both simulations, initial values of the state vector **x**(0) were sampled from a uniform distribution 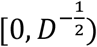. Initial values in the momentum vector **r**(0) were sampled from a standard Gaussian distribution.

### Simulation methods of replay of the linear track (Figure 4)

We assumed a linear track on which the position is indicated by a scalar *ψ* ∈ [0,1). We assumed that there were *D* = 100 place cells on the track, and *n*-th place cell had a Gaussian place field

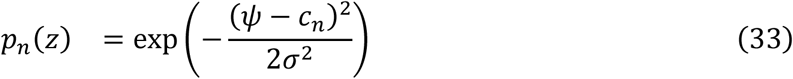

where the center and width of place fields are given as 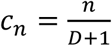 and *σ* = 0.2. We created memory patterns as **μ**_*i*_ = (*p*_0_(*ψ*_*i*_), …, *p*_*D*_ (*ψ*_*i*_))^*T*^ (*i* = 1, …, *M*) where 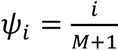. Then, memory patterns were normalized such that ∥**μ**_*i*_∥^2^ = 1.

Using these memory patterns, we simulated the momentum Hopfield model in the CA3-CA1 form by leap-frog integration (Eq. (22)-(25)). The state vector was initialized as **x**(0) = **μ**_0_ + (**μ**_0_ − **μ**_1_) to generate momentum from **μ**_0_ to **μ**_1_, which was transformed to **v**(0) = **Mx**(0) + 0.5. The constant 0.5 was added to initiate synchronized oscillation of **v**(*t*). The momentum **r**(0) was initialized by zero. Parameters were *β* = 30 and *ϵ* = 0.01.

Furthermore, for Bayesian place decoding, we created population activity vectors of binned positions as **ν**_*k*_ = (*p*_0_(*ψ*_*k*_), …, *p*_*D*_ (*ψ*_*k*_))^*T*^ (*k* = 1, …, *N*_bin_) where 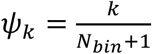. All **ν**_*k*_ were normalized such that ∥**ν**_*k*_∥^2^ = 1. The number of spatial bins was *N*_*bin*_ = 50.

### Simulation methods of replay of the 2-D field (Figure 5)

We assumed a square 2-D field in which the position is indicated by a Cartesian coordinate **ψ** = *ψ*_*x*_, *ψ*_*y*_ (*ψ*_*x*_, *ψ*_*y*_ ∈ [0,1)). We assumed that there were *D* place cells in the field, and *n*-th place cell had a Gaussian place field

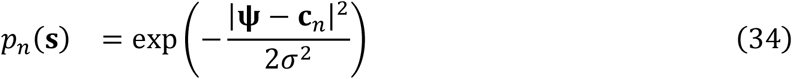

where *σ* = 0.1. The center of place fields **c**_*n*_ were randomly sampled in the field. We created memory patterns as **μ**_*i*_ = (*p*_0_(**ψ**_*i*_), …, *p*_*D*_(**ψ**_*i*_))^*T*^ (*i* = 1, …, *M*) where **ψ**_*i*_ were sampled independently from **c**_*n*_. Then, memory patterns were normalized such that ∥**μ**_*i*_∥^2^ = 1.

Using these memory patterns, we simulated the momentum Hopfield model by leap-frog integration in the CA3-CA1 form (Eq. (22)-(25). In the simulation, we resampled **v**(*t*) and **r**(*t*) every 10 unit time (*t* mod 10 = 0) and regarded each time period between resampling as an independent “replay event”. The vector **x**(0) was initialized by 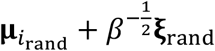 where *i*_rand_ is random interger in *i* = 1, …, *M* and **ξ** _rand_ is a sample from a *D*-dimensional standard Gaussian. Then, we initialized **v**(0) = **Mx**(0). The momentum was sampled from a standard Gaussian distribution. Parameters were *D* = *M* = 1000, *β* = 30 and *ϵ* = 0.01.

As in the simulation of the linear track, for Bayesian place decoding, we created population activity vectors of binned positions as **ν**_*k*_ = (*p*_0_(**ψ**_*k*_), …, *p*_*D*_(**ψ**_*k*_))^*T*^ (*k* = 1, …, *N*_bin_). Positions of spatial bins **ψ**_*k*_ were given by a 50 × 50 square lattice (*N*_*bin*_ = 2500). All **ν**_*k*_ were normalized such that ∥**ν**_*k*_∥^2^ = 1.

We performed Bayesian place decoding, and obtained a replay trajectory in each event **ψ**_decode_(*t*) by calculating positions with the maximum decoded probabilities at each time step. We calculated means of Euclidian distances of two positions of replay trajectories ‖**ψ**_decode_(*t*) − **ψ**_decode_(*t* + *Δt*)‖^2^ for *Δt* = 1, 1.1, 1.2, …, 3 and plotted log-log plot of mean traveled distances and time differences *Δ*t.

### Adding noise in the model (Figure 6)

To produce Brownian-motion-like replay by the model, we added noise to either the internal activation *v*_*i*_(*t*) or the pattern **μ**_*i*_. We introduced noise to *v*_*i*_(*t*) in the leap-frog integration as

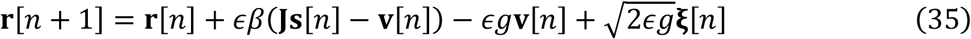

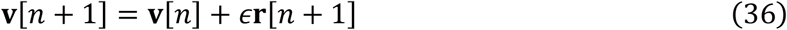

where **ξ**[*n*] is noise sampled from standard Gaussian. We generated noisy patterns as **μ**_*i*_ ← relu(**μ**_*i*_ + *ϵ* **ξ**_*i*_) where **ξ**_*i*_ is noise sampled from standard Gaussian and relu(*x*) is a rectified linear function. Adding noise was performed before normalization of the pattern. In both cases, the strength of noise was set as *g* = 0.05.

### Simulation methods of sampling of 2-D Gaussian mixture (Figure 7)

We tested two settings: bias-modulation and norm-modulation.

In the bias-modulation setting, we sampled elements in memory pattern vectors **μ**_*i*_ from uniform distribution [−1,1). We also randomly determined the mixing ratio as 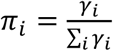 where *γ*_*i*_ is uniform random values in [0,1). Then, bias parameters were given as 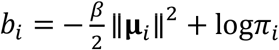.

In the norm-modulation setting, we sampled elements in memory pattern vectors from a standard Gaussian distribution. We also uniformly sampled values *γ*_*i*_ from [0.9,1), and normalized memory patterns such that |**μ**_*i*_|^2^ = *γ*_*i*_

Using 10 memory patterns generated as described above, we simulated the momentum Hopfield model in the original form by leap-frog integration (Eq. (20)(21)) for 10^6^ time steps. We note that, in the bias-modulation setting, bias parameters were introduced in the model (**x** ⋅ **μ**_*i*_ was changed to **x** ⋅ **μ**_*i*_ + *b*_*i*_ in Eq. (20)(21)). In the simulation, the momentum **r**(*t*) was resampled from a standard Gaussian distribution every 200 time steps. Parameters were *β* = 30 and *ϵ* = 0.05.

In both settings, recall of *i*-th memory was counted by ∑_*t*_ *s*_*i*_(*t*), and recall frequency was obtained by normalizing counts.

### Simulation methods of biased memory recall in 2-D field (Figure 7)

Although the procedure of the simulation was basically same with the simulation of 2-D replay, memory patterns were sampled at 20 x 20 grid positions in the 2-D field (not random positions). Heatmap in the result corresponds to recall frequency of these memories in the 20 x 20 grid. Furthermore, we multiplied 1.05 to memory patterns **μ**_*i*_ representing positions in a specific region (*ψ*_*x*_ < 0.4 and *ψ*_*y*_ < 0.4).

### Hippocampal network model for prioritized experience replay (Figure 8)

Based on the momentum Hopfield model in the CA3-CA1 form, we derived the hippocampal network model in which we can arbitrarily control sampling frequencies. We consider the network in which sampling frequencies follow

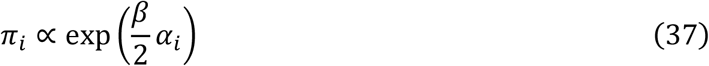

where parameters *α*_*i*_ are arbitrary constants that control sampling. Such sampling is obtained by embedding memory pattern 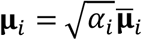 where 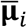 is unit-norm pattern vectors. Then, we obtain the model as follows:

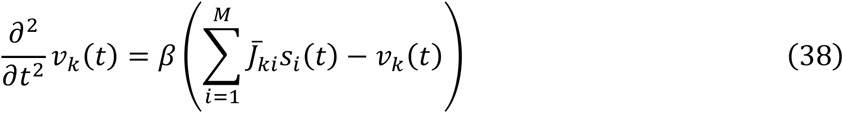

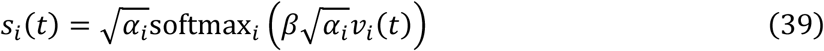

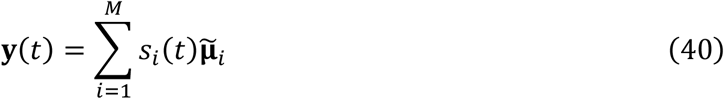

where 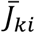 is normalized weight matrix 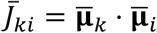.

### Setting of reinforcement learning for the spatial navigation task (Figure 8)

In the simulation of prioritized experience replay, we assigned a memory pattern to each episode for TD learning *e* = (*s, s*_next_, *a, r*) (the memory pattern **μ**_*i*_ corresponds to *i*-th episode *e*_*i*_). The current state *s* is a position in the 20 × 20 discretized 2-D space. The action *a*_*t*_ is a movement to one of four directions (North, East, South, West), and *s*_next_ corresponds to the position after the movement from *s*. Reward *r* is 1 if the agent gets to the goal (the current position *s* is the goal); otherwise *r* = 0. In the simulation, instead of the exploration in the 2-D space, we iteratively sampled episodes from *ε* which is a set of all possible combinations of (*s, s*_next_, *a, r*) in the space (experience replay). Using sampled episodes, the agent learned Q-value functions *Q*(*s, a*) by Q learning as

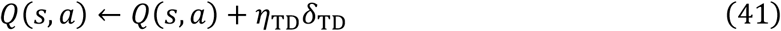

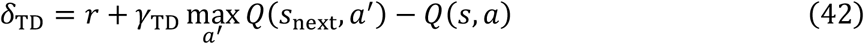

where *η*_*TD*_ = 0.5 and *γ*_*TD*_ = 0.9. Initial values of *Q*(*s, a*) were uniformly sampled from [0, 0,1). Sampling and learning were performed 10^5^ times in each condition. We periodically recorded Q functions during learning, and evaluated the performance of spatial navigation from the start to the goal by a *ϵ*-greedy policy (*ϵ* = 0.1). We repeated 100 trials of navigation for each Q function, and calculated the average number of time steps required for getting to the goal. We terminated the navigation if the agent did not get to the goal within 200 time steps.

### Simulation methods for experience replay (Figure 8)

We tested four methods to sample episodes: “TD-bias”, “reward-bias”, “unbiased”, and “random”. The “random” method is uniform random sampling of all possible episodes. In other sampling methods, we simulated the momentum Hopfield model in the CA3-CA1 form by leap-frog integration (Eq. (22-25)), and we sampled an index of the recalled memory as *i*_max_ = argmax_*i*_ *s*_*i*_(*t*) every 5 time steps of the leap-frog integration. Then, an episode corresponding to the sampled memory 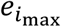 is replayed and TD learning was performed. In the simulation, the momentum **r**(*t*) was resampled from a standard Gaussian distribution every 100 samples. Parameters were *β* = 30 and *ϵ* = 0.05.

In the “TD-bias” method, the parameter *α*_*i*_ for the replayed memory pattern was substituted by the absolute TD error as *α*_*i*_ = *a*_TD_|*δ*_TD_| + *b*_TD_ every time TD learning was performed (*a*_TD_ = 2, *b*_TD_ = 0.5). In the “reward-bias” method, *α*_*i*_ is determined by proximity of the corresponding episode (*e*_*i*_ to rewarded episodes (a set of episodes in which the current state is the goal *ε*_*g*_ as

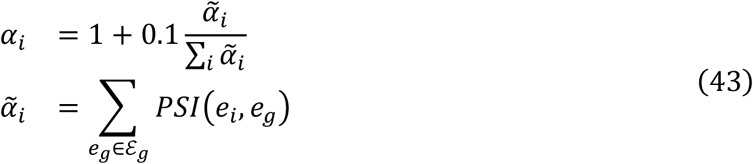

where proximity is evaluated by positive successor information *PSI*(*e, e*′) described below. In the “unbiased” method, *α*_*i*_ was fixed to 1.

We created memory patterns for episodes using disentangled successor information with sparsity (DSI-sparse) ^45^, which is a hippocampal representation model based on successor representation (SR) ^46,47^. Assuming that we have SR for episodes *SR*(*e, e*′) and stationary probability distribution *P*(*e*), SI and PSI (positive successor information) are defined as

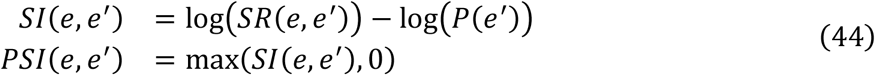

We performed dimension reduction of PSI as **x**(*e*) ⋅ **w**(*e*′) ≈ *PSI*(*e, e*′) where **x**(*e*) and **w**(*e*) were 50-dimensional non-negative vectors in this simulation. This is performed by optimization of the following objective function by Nesterov’s accelerated gradient descent.

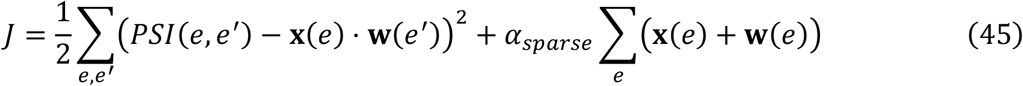

We rectified **x**(*e*) and **w**(*e*) every iteration. Learning rate was 0.05, the parameter *α*_*sparse*_ was 10^−4^. We created memory patterns for episodes by normalizing **x**(*e*).

In the calculation above, *SR*(*e, e*′) and *P*(*e*′) were calculated from an adjacency matrix of episodes **A**^ep^. In **A**^ep^, each element 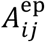 is 1 if there is possibility of the transition from *e*_*i*_ and *e*_*j*_ (i.e. *s*_next_ in *e*_*i*_ is same with *s* in *e*_*j*_); otherwise 0. Assuming random walk, we obtained the transition matrix for episodes **T**^ep^ by 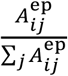. Then, *SR*(*e, e*′) and *P*(*e*′) were obtained as following.

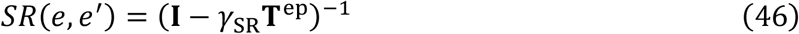

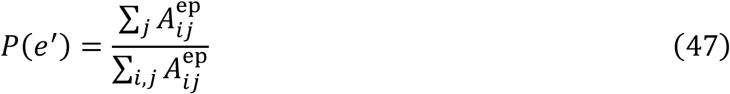

where *γ*_*SR*_ was 0.9.

## Acknowledgements

This research was partially supported by JST CREST JPMJCR23N2 and JST PRESTO JPMJPR2519.

