## Supplementary figures and legends of videos for "Efficient memory sampling by hippocampal attractor dynamics with intrinsic oscillation"

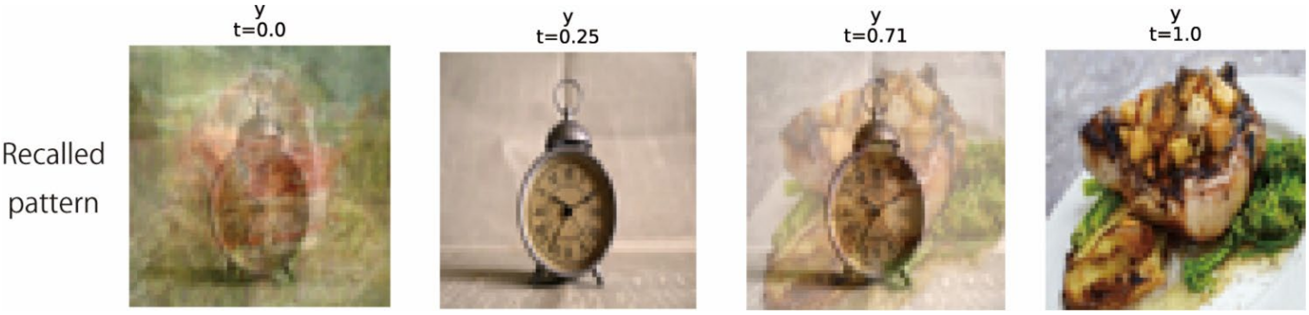


**Supplementary Figure 1:** Image retrieval by momentum Hopfield model in the CA3-CA1 form.

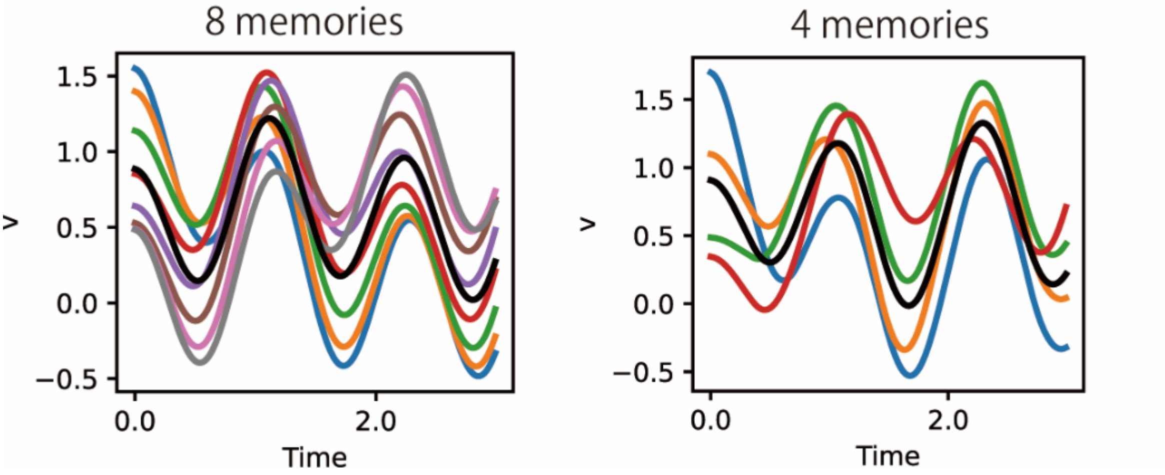


**Supplementary Figure 2:** Oscillation of $v_k(t)$ in the simulation of replay on the linear track.

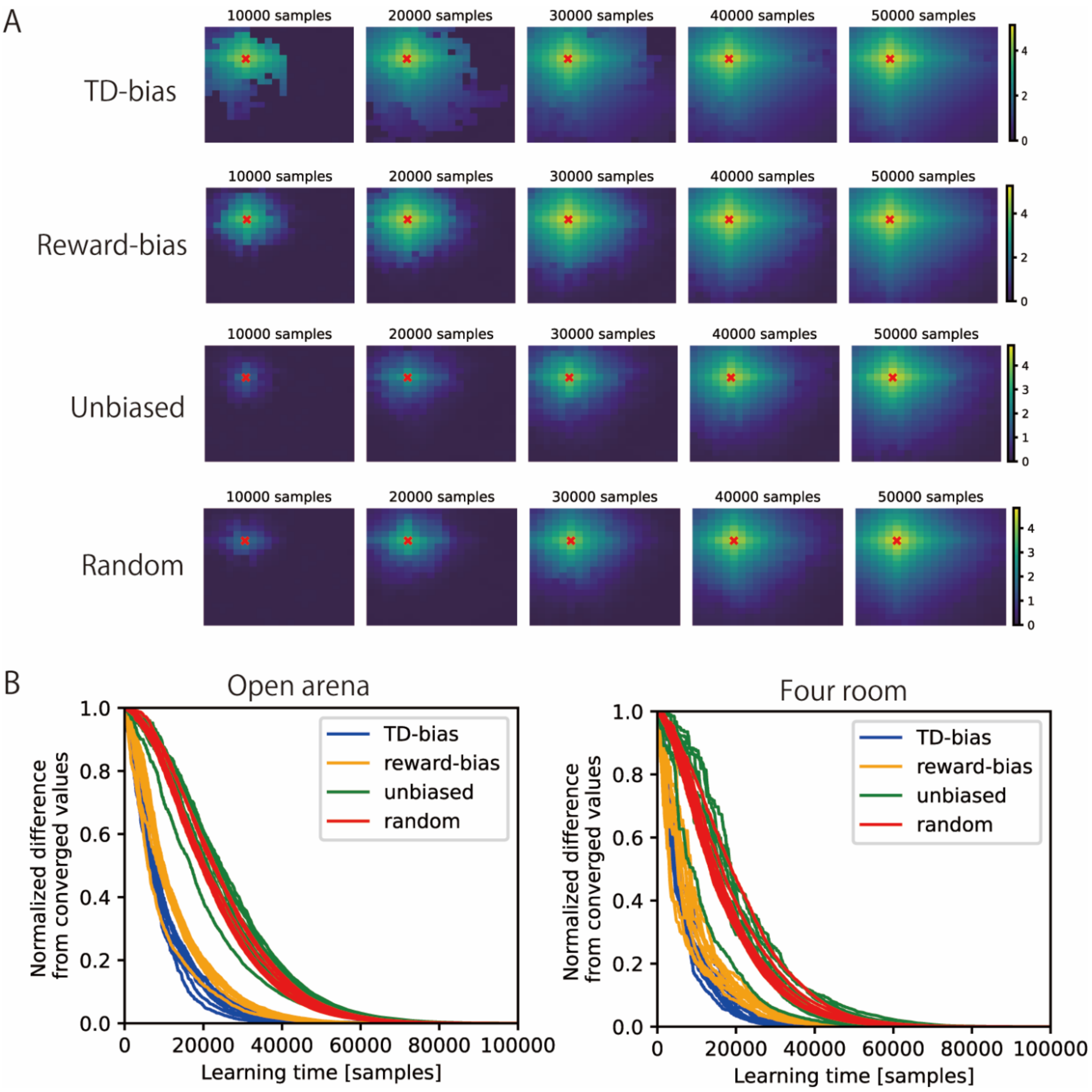


**Supplementary Figure 3:** Comparison of value learning by four replay methods. (A) Heatmaps of value functions in the environment learned by four replay methods. (B) Comparison of convergence speeds of value functions (normalized differences from the final values) based on various replay methods (10 independent simulations for each setting).

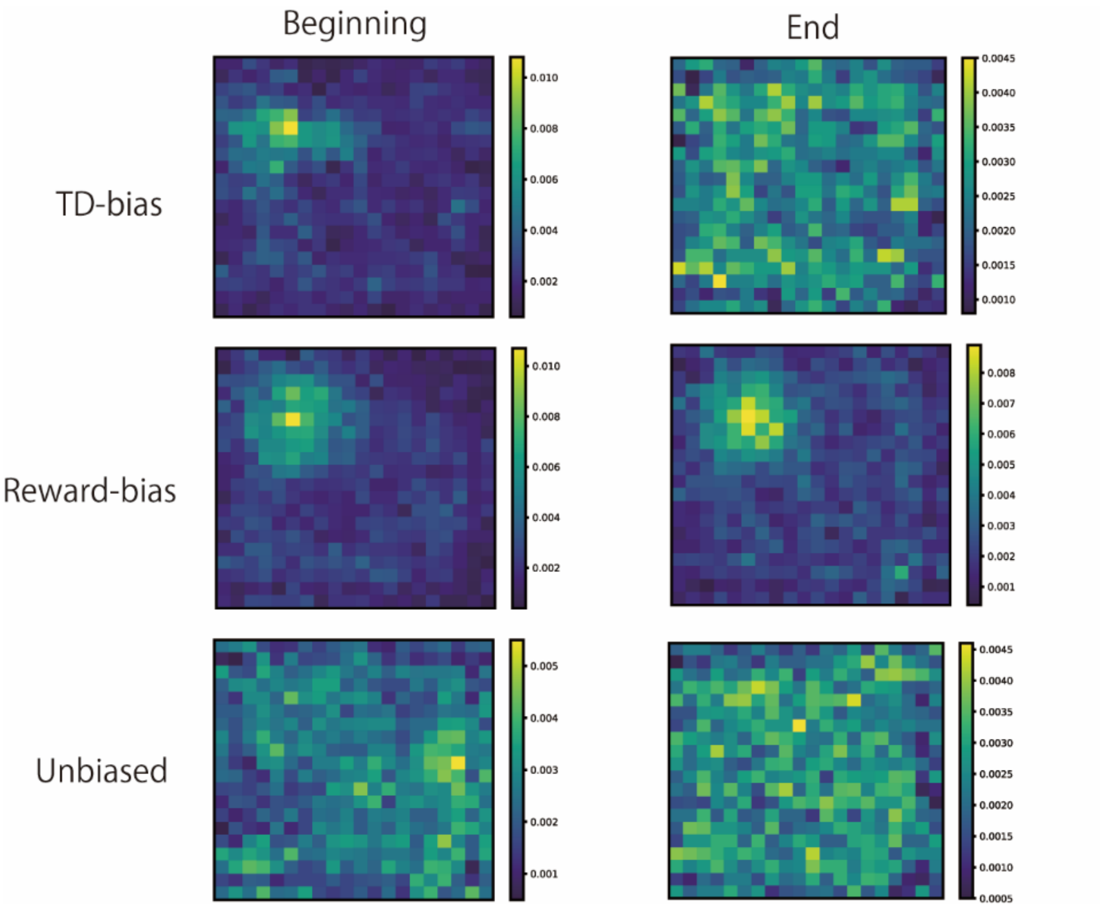


**Supplementary Figure 4:** Replay frequencies of each position in the open-arena setting by various replay methods (TD-bias, reward-bias, unbiased). Left: frequencies in 10,000 samples at the beginning of learning. Right: frequencies in 10,000 samples at the end of learning.

**Legends of supplementary videos**

Supplementary video 1: Time evolution of x(t) and y(t) in image retrieval by the momentum Hopfield model (corresponding to Figure 1).

Supplementafy video 2: Decoded probability distributions (heatmap) and replay trajectories (red lines) in the simulation of 2-D place-cell replay by the momentum Hopfield model (corresponding to Figure 5).
